# 4D Mesoscale liquid model of nucleus resolves chromatin’s radial organization

**DOI:** 10.1101/2023.09.21.558819

**Authors:** Rabia Laghmach, Michele Di Pierro, Davit A. Potoyan

## Abstract

Recent chromatin capture, imaging techniques, and polymer modeling advancements have dramatically enhanced our quantitative understanding of chromosomal folding. However, the dynamism inherent in genome architectures due to physical and biochemical forces and their impact on nuclear architecture and cellular functions remains elusive. While imaging techniques capable of probing the physical properties of chromatin in 4D are growing, there is a conspicuous lack of physics-based computational tools appropriate for revealing the underlying forces that shape nuclear architecture and dynamics. To this end, we have developed a multi-phase liquid model of the nucleus, which can resolve chromosomal territories, compartments, and nuclear lamina using a physics-based and data-informed free energy function. The model enables rapid hypothesis-driven prototyping of nuclear dynamics in 4D, thereby facilitating comparison with whole nucleus imaging experiments. As an application, we model the *Drosophila* nucleus spanning the interphase and map phase diagram of nuclear morphologies. We shed light on the interplay of adhesive and cohesive interactions within the nucleus, giving rise to distinct radial organization seen in conventional, inverted, and senescent nuclear architectures. The results also show the highly dynamic nature of the radial organization, the disruption of which leads to significant variability in domain coarsening dynamics and, consequently, variability of chromatin architecture. The model also highlights the impact of oblate nuclear geometry and heterochromatin sub-type interactions on the global chromatin architecture and local asymmetry of chromatin compartments.

## I. INTRODUCTION

Advances in genomics, computer simulations, and high-resolution microscopy have provided unprecedented insights into chromatin folding and its functional implications [1, 2]. Three distinct mechanisms responsible for shaping nuclear architectures have emerged [3]; loop extrusion, phase separation, and chromatin anchoring to the nuclear envelope. The physical interactions driving these mechanisms are lengthwise compaction, chromatin loci cohesion, and chromatin loci adhesion to the nuclear membrane. The lengthwise compaction [4–7] originates from an interplay of equilibrium protein binding and non-equilibrium machinery that loop distal regions of chromosomes by forming local structures referred to as Topologically associating domains (TADs). Cohesive interactions [8, 9], which lead to phase-separation (PS) and of chromosome regions originate from the presence of epigenetically distinct A/B chromatin loci, which drive the formation of heterochromatin (HC) and Euchromatin domains (EC), respectively. The origin of adhesive interactions between chromatin and the nuclear membrane [10] is mediated by filaments, collectively known as nuclear lamins. Lamins interact, directly or indirectly, with heterochromatin, resulting in preferential anchoring of heterochromatin to the nuclear membrane. Genomic regions that have the ability to anchor to nuclear envelope can be identified in experiments and are referred to as Lamineassociated domains (LADs).

Chromatin architecture lives in an inherently active and stochastic environment surrounded by numerous proteins and RNA, which constantly remodel and reorganize chromatin architecture [11–13]. At the same time, architectural dynamical patterns of chromatin organization are not random but instead are tightly coupled to the embryonic developmental timetable and cell cycle. Numerous experiments report on the radial organization of the genome’s physical features, including GC content gradients, transcriptional activity, and heterochromatin distribution relative to nuclear origin [14–18]. For instance, in healthy cells, nuclei adopt a conventional architecture with a distinct radial preference of heterochromatin regions towards nuclear envelope [19]. Conversely, heterochromatin gradually shifts toward the center in the senescent cells, often forming a few larger clusters [20]. During differentiation, heterochromatin may detach from lamina altogether, forming the so-called inverted nuclei, which are common in the rods of nocturnal mammals [19]. Cancer progression is likewise accompanied by a massive reorganization of heterochromatin regions near lamina [20, 21]. Several fundamental questions remain unanswered regarding the role of lamina loss and its impact on chromatin organization during senescence and various diseases. The increasing sophistication in live nuclei imaging, 3D microscopy, and spatial transcriptomics [22–24] are now generating data that could benefit from mechanistic 3D nucleus modeling approaches. These advances open road for mechanistic modeling of nuclei on mesoscopic scales, paving the road for uncovering the link between the mechanobiology of the nucleus and its transcriptional activities.

To this end, we have developed a mesoscale model of the nucleus termed the Mesosclale Liquid Model of Nuclear Dynamics in 4D (MELON-4D). The model uses a physics-based and data-informed free energy function to describe the dynamic evolution of chromosomal territories, compartments, and nuclear lamina. The physics-based part of the free energy function is motivated by experimental features of the eukaryotic genome learned from Hi-C and imaging experiments. Specifically, the model accounts for the three fundamental driving forces of the chromosomal organization by explicitly modeling cohesive interactions between liquid chromatin phases, adhesive interaction with the lamina, and differential mobilities of euchromatin and heterochromatin due to different degrees of local compaction. At the same time, the model is not fitted to data that has allowed us to study in detail the interplay of forces underlying key characteristics of 3D nuclear architectures, including radial distribution of heterochromatin, domain-domain distances, shapes, and volumes of domains. Finally, we note that overall, our results regarding the radial organization and its role as a gatekeeper of architectural variability are in harmony with the findings of recent computational models of *Drosophila* genome [25].

## II. MELON-4D

Here we describe the MELON-4D, a model for describing the mesoscale motions of chromatin domains in 3D nuclear geometries. This model builds upon our previous phase-field approach, which focused on chromatin-type patterning in a fixed 2D geometry [26, 27]. The previous model successfully recapitulated several experimentally observed phenomena, including phase-separation-driven inter-chromosomal coherent motions, chromatin patterning in aging and normal nucleus, and activity-enhanced droplet fusion dynamics. However, the 2D model of the nucleus had severe shortcomings as it surfaces tension and fusion dynamics of chromatin compartments in 3D space. Below we summarize the multiphase field formulation used in the MELON-4D model by highlighting key improvements relative to our previous approach. In the MELON-4D, a set of phase fields variables ***φ***(**r**, *t*) = {{*φ*_*i*_(**r**, *t*) }_*i*=1,…,*N*_}, and ***ψ***(**r**, *t*) = {{*ψ*_*j*_(**r**, *t*)} _*j*=1,2_} are introduced as non-conserved order parameters to describe the shape variation of various compartments within the nucleus that are corresponding to N-chromosomal territories, and three types of chromatic regions of chromosomes which are in euchromatin (EC) and facultative/constitutive heterochromatin forms (fHC and cHC). The phase-field variables *φ*_*i*_(**r**, *t*) and *ψ*_*j*_(**r**, *t*) vary smoothly across their interfaces profile between two values that it is 1 inside its compartment and 0 elsewhere. The geometry of the nucleus is defined through an auxiliary order parameter *η*(**r**), which takes 0 inside the nucleus and 1 outside and varies smoothly between these two values through the interfacial region. This region represents the nuclear envelope whose position is given by the iso-contour η = 1/2. The field η(r) is used as an indicator function independent of time to model a fixed nucleus with volume *V*_*N*_. The nuclear envelope is assumed here at the equilibrium state during the interphase of the cell cycle, which can be represented by a tanh-like profile of η(r). We simulated the oblate nuclear shape typical of eukaryotic nuclei during interphase by the use of the following expression: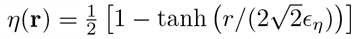; where 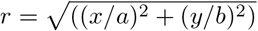 is the distance from the center of the physical compartment Ω, whereas *a* and *b* are the semi-major and semiminor<u>a</u>xes. The width of the nuclear envelope is given by 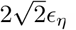.

The dynamics of the chromatin compartmentalization patterns are derived from the free energy functional ℱ [***φ, ψ***], which describes the intra-nuclear phase separation of chromatin subtypes. The evolution of the phasefield variables {***φ, ψ***} are governed by the Allen-Cahn equations:

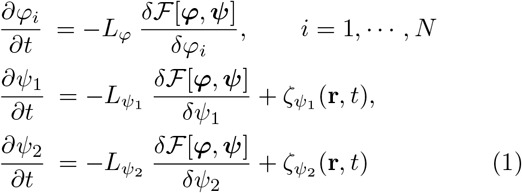

where 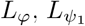 and 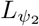 are mobility coefficients that are proportional to the relaxation time of different phase-field variables. We have chosen the mobility coefficients to match the magnitudes of *in vivo* measurements of euchromatin and heterochromatin diffusion coefficients [28] The terms 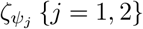 in Eq. (1) account for the fluctuations at the boundaries of EC/fHC and EC/cHC islands due to finite size nature of the droplets. The fluctuations are modeled as white noise: 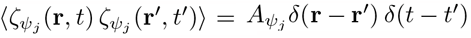 where the amplitude of the noise is given by: 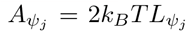 . The amplitude of noise *A*_*p*_ sets the “effective temperature” *T*_*eff*_ of the nucleus [29], which can be taken as a measure of ATP activity in comparisons with the experiment [30, 31].

The phase-field description of the nucleus in the present work is best described in terms of experimentally motivated constraints on shapes and sizes of the nucleus and chromosomal territories, which are supplemented by the physics-based terms accounting for polymer intermingling diffusive segment motion within and between chromosomal territories. The primary driving forces for emergent nuclear architecture and dynamics are derived from the microphase separation of heterochromatin sub-types, the surface tension of chromatin droplets, and differential affinity for chromatin-lamina interactions. Chromatin types’ volume and surface constraints are imposed to capture chromosomal and nuclear boundaries. The full free energy is functional of the nuclear chromatin ℱ [***φ, ψ***], which we minimize in Eq. (1) to get the evolution equations of the nuclear structures can be split into two energy functional contributions *F*_*B*_ and *F*_*I*_ . The GinzburgLandau free energy functional *F*_*B*_ describes the coexistence of two phases associated with each phase-field variable completed by volume constraints terms to ensure the shapes change given by:

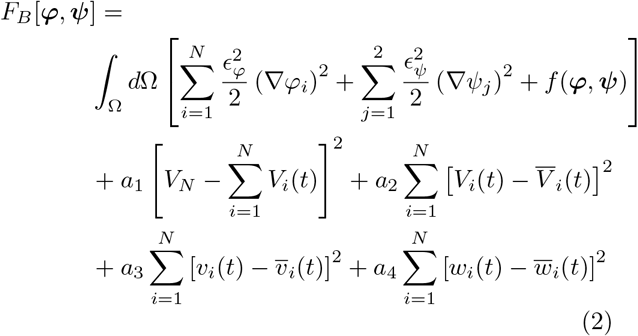

where *f* (***φ, ψ***) is the bulk free energy contribution for multi-phase field variables, and gradients term accounting for the presence of different interfaces in the system and contributing to the interfacial energies. The gradient parameters *ϵ*_*φ*_ and *ϵ*_*ψ*_ control the thickness of the interface profile of ***φ*** and ***ψ***, respectively. For the bulk free energy density, we use a multi-well potential expressed as: 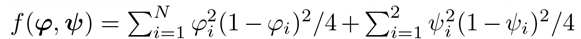. The terms are proportional to *a*_*i*_ account for volume constraints required to enforce the volume of the chromosomal territories at their prescribed values 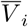, the facultative heterochromatin at 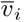 and the constitutive heterochromatin at 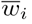. The parameters *a*_1_, *a*_2_, *a*_3_ and *a*_4_ are positive coefficients that control the thermodynamic driving forces of coarsening processes of different compartments present in the nucleus. The volumes of the *i*chromosomal territory *V*_*i*_(*t*), facultative and constitutive heterochromatin compartments within each chromosome *v*_*i*_(*t*) and *w*_*i*_(*t*) are defined as the spatial integral over the physical compartment Ω of their interface profiles given by the associated phase-field variables *φ*_*i*_(**r**, *t*), *ψ*_1_(**r**, *t*) and *ψ*_2_(**r**, *t*). Using the usual phase-field approximation of the volume could change the coexistence phase values defined by the phase-field variables between 0 and 1.We thus used an interpolation function, defined as 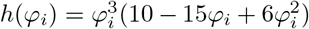, for approximating the volumes of different compartments of the nucleus while keeping the position of the local free energy minima at the coexistence phase values. The following expressions approximate the volumes of these compartments: *V*_*i*_(*t*) = *d*Ω *h*(*ψ*_1_)*h*(*φ*_*i*_); and *w*_*i*_(*t*) = *d*Ω *h*(*ψ*_2_)*h*(*φ*_*i*_). Next, we define the free energy functional *F*_*I*_, which accounts for the geometrical constraints on the nucleus, excluded volume interactions between different chromatin compartments within the nucleus, and the interaction between heterochromatic subtypes with the nuclear envelope. The functional *F*_*I*_ is expressed as:

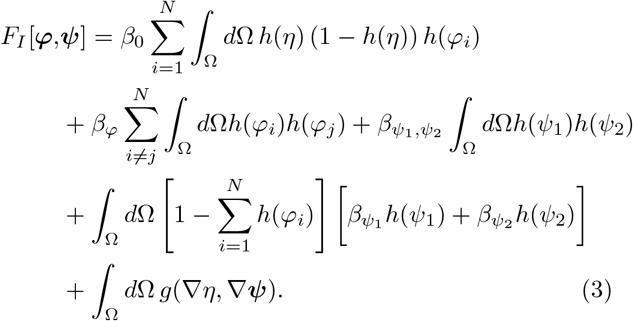

The first term in *F*_*I*_ corresponds to the energy penalty reflecting the geometrical constraint on the nuclear volume required to restrain nuclear components’ motion inside the nucleus. The other terms represent the excluded volume interactions between chromosome territories, fHC-cHC regions interactions, and HC/EC regions mixing affinity, with the strengths of the interactions described by the parameters 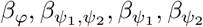, respectively. The last term represents the interaction of the fHC and cHC with the nuclear envelope through a *g* function which represents the local Lamina-interaction energy contribution to the free energy functional:

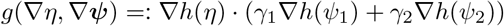

where *γ*_1_ and *γ*_2_ are two positive parameters controlling the binding affinity of heterochromatin types to the nuclear lamina.

### Parametrization of the MELON-4D

To numerically solve the set of evolution equations of the phase-field variables resulting from the free energy minimization (1), we use the finite element method combined with the preconditioned Jacobian Free Newton Krylov approach (JFNK). The model has been implemented in MOOSE finite element C++ library [32, 33], which is built using high-performing computational libraries MPI, LibMesh, and PETSc needed for solving non-linear partial differential equations [33].

The computational domain used here is set as Ω = [0, *L*_*x*_] × [0, *L*_*y*_] × [0, *L*_*z*_], with *L*_*x*_ = 6 *μm, L*_*y*_ = 9 *μm, L*_*z*_ = 3*μm*, and we meshed this domain to generate a fine mesh made up of 180 ×225 ×90 elements. The time step used for time integration is set to 0.04 in dimensionless units, which is chosen to ensure the numerical stability of all simulations. The nuclear shape is maintained in an elliptical shape in which the semi-axes are fixed to *a* = 2.5 *μm, b* = 4 *μm*, and *c* = 1.2 *μm*; thus, the nuclear volume is: 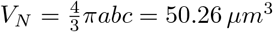 consistent with empirical measurements of *Drosophila* nucleus during interphase [34]. The mobility of chromosomal and heterochromatin compartments are set to be equal 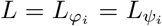 in all the simulations. This parameter fixes the compartment’s interface relaxation time: *τ* ∝ *L*^−1^. The interface relaxation time *τ* is used here to set the unit timescale for the kinetics of phase transitions. The length scale is fixed at *l* = 1 *μm*. We set *τ* = 0.005 *s* for all simulations. The diffusion coefficient can be expressed as 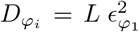 for chromosomal territories, 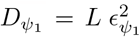 for the heterochromatin and 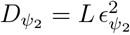 for chromocenter. For all the simulations, the diffusion coefficients are fixed as 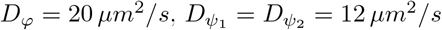. These values of diffusion coefficients are motivated by experimental measurements of euchromatin/heterochromatin mobility in live cells [28]. The interaction coefficient between chromosomal territories and the nuclear envelope is strong enough to maintain them inside the nucleus by setting *β*_0_ = 16.7. To ensure well-separated chromosome territories, we set the chromosome-chromosome interaction parameter with strong interaction like the one in [26] *β*_*φ*_ = 40. The heterochromatin-heterochromatin interaction parameter is taken as 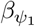 to 0.1 for all simulations, and chromo-chromocenter and hetero-chromocenter interaction parameters 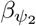 and 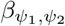 are varied. The interaction parameters between heterochromatin and nuclear lamina, *γ*, is varied from 0 to 1.1 to evaluate the effects of competition between binding energies and chromatin compartment interactions. The remaining model’s parameters are fixed to enable dynamics comparable to chromatin diffusion coefficients: *a*_1_ = 0.16 and *a*_2_ = *a*_3_ = *a*_4_ = 2. The fluctuation amplitude of euchromatin-heterochromatin and euchromatin-chromocenter interfaces *A*_1_ and *A*_2_ are fixed at 5. While the nuclear shape, size, chromosome numbers, and diffusion coefficients are calibrated after *Drosophila* nucleus, the model is sufficiently general for drawing broader conclusions about the chromatin structure and dynamics in eukaryotic nuclei.

## III. RESULTS

We model the *Drosophila* nucleus using an elliptical geometry containing eight chromosomal territories *N* = 8 (See Methods). Each chromosome is resolved at the level of chromatin types corresponding to epigenetically distinct euchromatin (EC) and heterochromatin (HC) compartments. In the following two sections, we look at architecture and the dynamic evolution of heterochromatin compartments formed with one-component and two-component nuclei corresponding to constitutive (cHC) and facultative (fHC).

In the MELON-4D, all phase-separated chromatin compartments are defined by phase-field variables tracking interfaces, volumes, and geometries of chromatin compartments. To evaluate how the interactions between chromatin types impact the organization and dynamics of compartments, we perform simulations of the 3D nucleus by varying the interaction strengths between chromosomal territories and different chromatin types. Note that the interactions between chromosomal territories govern the degree of intermingling between neighboring CTs and the global arrangement of chromosomes in the interphase nucleus. CT interactions are long-ranged and correspond to the nucleus’s slowest timescale of chromatin motions. CTs interaction are controlled by the parameter *β*_*φ*_. Microphase separation and motion of chromatin compartments correspond to faster timescale motions in the nucleus. The parameter 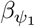 in the two-component chromatin model controls the attraction strength between EC-HC within individual CT. Higher values of 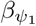 correspond to stronger attraction between chromatin types within CTs. Likewise, two parameters controlling attraction strengths between Euchromatin (EC) and heterochromatin (HC) subtypes are 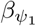 and 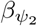corresponding to constitutive and facultative heterochromatin respectively. The strength of interaction between the HC subtypes is given by 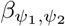 . A higher value of 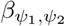 indicates weak attraction between the subtypes of heterochromatin. Thus, increasing the value of this parameter could lead to phase-separated heterochromatin compartments.

### A. Nuclear organization resolved with a two-component chromatin model

Here we consider two interacting chromatin types, Euchromatin (EC) and Heterochromatin (HC). The dynamics and morphology of heterochromatin compartments in the long time limit are controlled by chromatin-type interactions within chromosomes and the degree of intermingling between neighboring chromosome territories.

First, we carried out simulations with fixed strength of CTs interaction. Predictably, for stronger chromatin type-to-type attraction, we find disconnected heterochromatin droplets in individual chromosomes (Fig. 2A). For weaker chromatin type-to-type attraction, we find connected heterochromatin droplets of different chromosomes residing in the interior of the nucleus (Fig. 2A). Furthermore, weaker chromatin type-to-type attraction revealed a more pronounced clustering of heterochromatin droplets, resulting in two large compartments localized within the interior of the nucleus (Fig. 2A). The nuclear structure obtained with stronger chromatin typeto-type attraction showed that heterochromatin droplets formed within chromosomes maintained their positions in the center of each CT and are surrounded by euchromatin. Thus an increase in chromatin type-to-type attraction drives the localization of heterochromatin compartments within the chromosomal territory. Interestingly, in our previous work using a two-dimensional model of the nucleus [26], the emerged nuclear structure with stronger chromatin type-to-type attraction has exhibited fewer heterochromatin droplets clustering in the interior of the nucleus. This result shows the importance of 3D motions even for oblate geometry, which more accurately accounts for spatially interacting chromatin types that regulate the movement and clustering of the formed heterochromatin droplets.

**FIG. 1.**
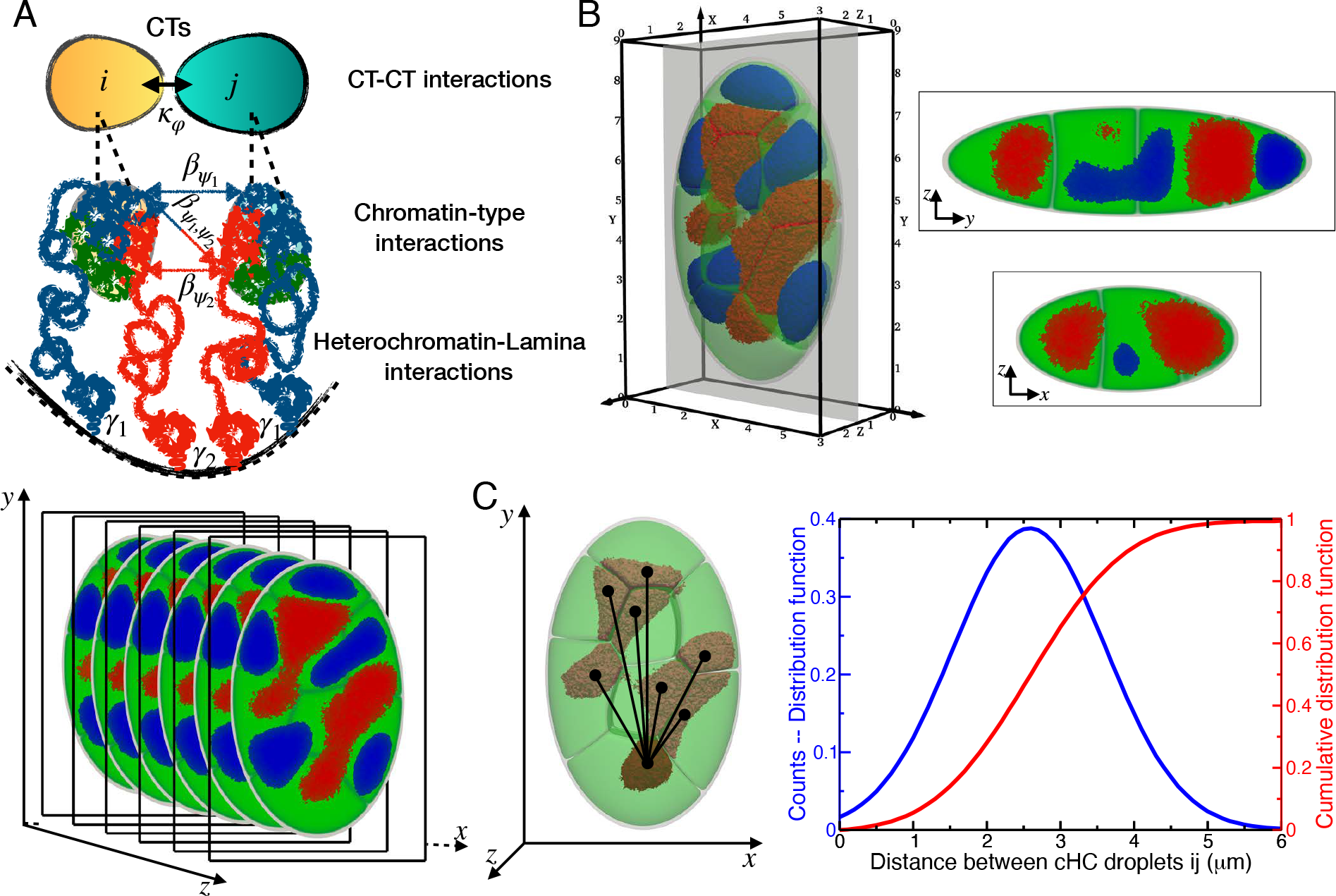
(A) 3D snapshots of a model *Drosophila* nucleus simulated using the MELON-4D framework resolving three chromatin types: euchromatin (green), facultative (blue), and constitutive heterochromatin (red). (B) A 2D slice snapshot of simulated nuclear architecture along the x-y, y-z, and x-z axes. (C) Distribution of distances between the heterochromatin compartment centroids from the *i*-th and j-th chromosomes (*j* ≠ *i*).

**FIG. 2.**
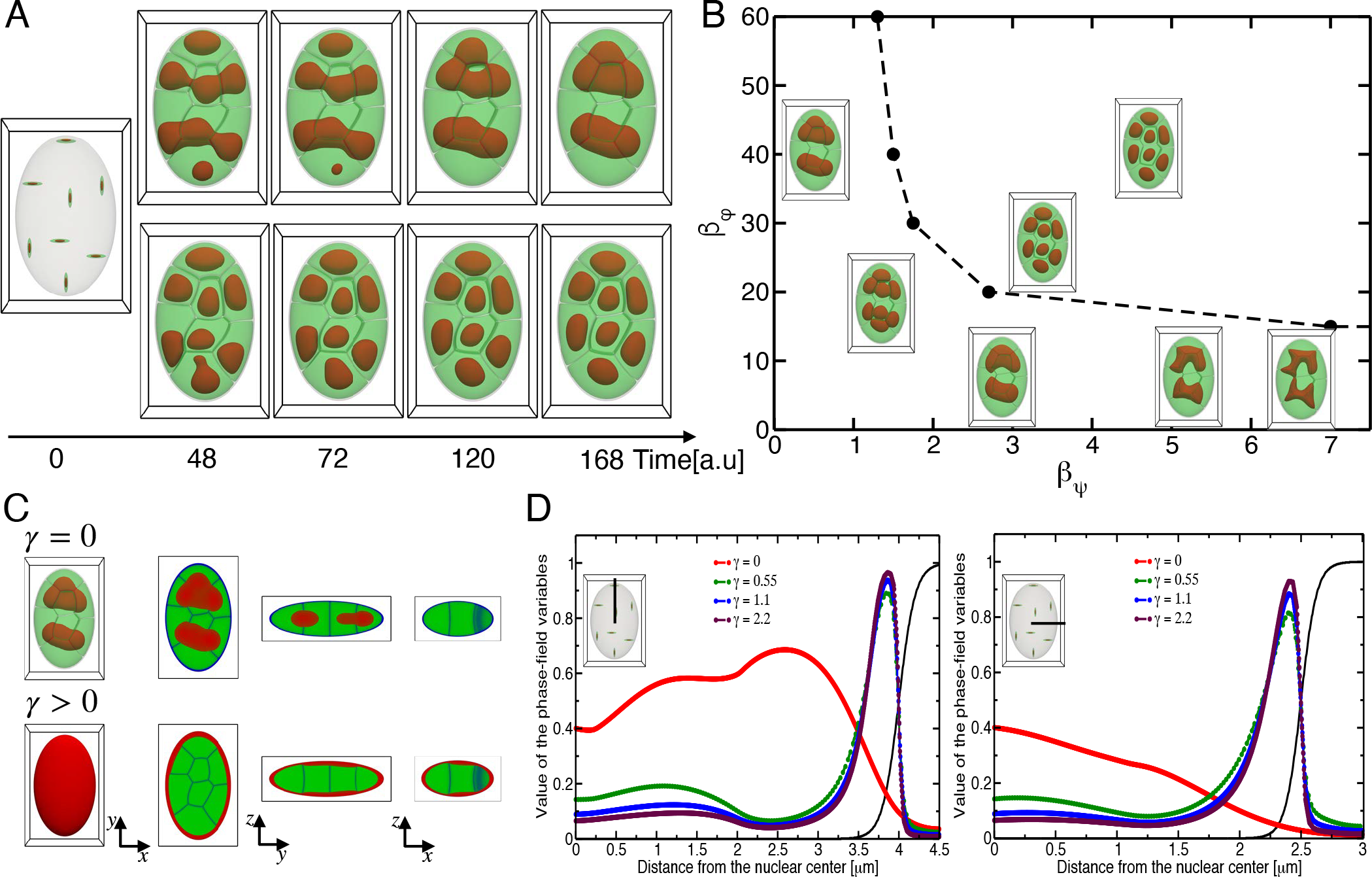
Temporal evolution of chromatin A/B patterns in 3D (A) Shown the effect of chromatin-type interactions on the degree of spatial compartmentalization of the nucleus. The top panel shows the emergent chromatin pattern driven by a weaker attraction between chromatin types within chromosomes and the bottom panel shows stronger attraction. (B) The phase diagram of nuclear chromatin architectures as a function of the two parameters controlling the strength of interactions between chromosomes and chromatin types. (C) Shown are 3D nuclear architectures and 2D slices corresponding to the absence (*γ* = 0) and presence (*γ >* 0) of adhesive lamina-heterochromatin interactions in the top and bottom panels, respectively. (D) Radial density profile of heterochromatin along the major and minor axis computed from the nuclear center as a function of adhesive lamina-heterochromatin interaction strength.

The interplay between CTs and chromatin type-totype interactions can span a wide spectrum of nuclear architectures. Therefore, we next vary the two parameters controlling the strengths of physical interactions between CTs and chromatin types in the nucleus. Results are presented as phase diagrams of nuclear architectures showing the connectivity between heterochromatin droplets of different chromosomes (Fig. 2B). We identify three distinct morphological regimes in the phase diagram.

The first morphological regime corresponds to stronger chromatin type-to-type but weaker CTs interactions, driving disconnecting heterochromatin droplets formed within individual chromosomes. The second regime corresponds to weaker chromatin-type and CTs interactions, which drive the formation of strongly connected heterochromatin droplets localized near the center of the nucleus. The last regime corresponds to stronger chromatin type-to-type and CTs interactions, which drive the formation of strongly connecting heterochromatin droplets across the boundaries of chromosome territories. We note that morphologies generated by modeling have been observed in experiments on eukaryotic nuclei in stages of cell life where the connection with lamina is severed, including embryonic growth of some species, inversion, and senescence [19, 35–37].

Having considered cohesive interactions between chromatin types in 3D nuclei, we next turn on the typespecific adhesive interactions between chromatin with lamina and analyze radial profiles and volumetric characteristics of heterochromatin-enriched domains. Radial profiles of Drosophila and other nuclei have been measured in recent whole genome experiments [36, 38, 39] and explored in the whole nuclei simulations of Drosophila [25] pointing out their role in generating robust non-random average global architectural patterns of heterochromatin which is relatively insensitive to type interactions. Experimentally, it is known that heterochromatin compartments are partially tethered to the nuclear lamina for healthy *Drosophila* and other eukaryotic nuclei [25, 38]. A preferential interaction between heterochromatin and the nuclear lamina could be sufficient to drive the motion of heterochromatin towards the nuclear periphery and lamina-anchoring of heterochromatin droplets.

In our two-component chromatin model with sufficiently adhesive interactions (Fig. 2CD), we are able to capture nuclear architectures with radial profiles resembling the experiments and whole nuclei simulations of Drosophila [25, 38]. First by looking at the global features of radial profiles in 3D nuclei, we find the importance of analyzing both major and minor access since some of the domain formations may be masked when taking a 2D slice-representation of nuclei which is done in experiments and 2D continuum models [26]. We found that the strength of the adhesive interaction, besides impacting chromatin 3D architecture, also has consequences on the dynamics of heterochromatin droplet formation around the nuclear envelope. That is, stronger heterochromatin adhesive interaction generates rapid quenching of heterochromatin dynamics, leading to differences in 2D radial profiles between the major and minor axes (Fig. 2CD). Overall the results demonstrate how the interplay of chromatin interactions could regulate chromatin dynamics within the nucleus and shape conventional and inverted nuclear architectures. Additionally, the 2D slice-based representation shows that chromatin distribution and variation in the size of heterochromatin compartments are spatially associated with the characteristic dimensions of the nucleus (Fig. 2C). Based on this observation, we hypothesize that the differences in lamina attachment rates can impact gene expression variability, which can be quantified by studying the variability of heterochromatin layer thicknesses near the lamina.

### B. Nuclear organization resolved with a three-component chromatin model

Here, we consider the three-component liquid chromatin model of the nucleus which can account for additional interactions taking place between chromatin subtypes, namely constitutive cHC and facultative fHC types of heterochromatin. Physical interactions between chromosomes CTs and all distinct pairs of chromatin subtypes (cHC, fHC, EC) now govern the motion of heterochromatic compartments through the chromosomal boundaries, resulting in distinct architectures.

To assess the sensitivity of the global radial order of the nucleus’ compartmentalization to chromatin subtype interactions, we performed 3D simulations by varying the interaction strength of 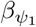 between cHC-EC and of 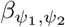 between cHC-fHC types of chromatin. Having shown that maintaining separated chromosomal territories with the possibility of overlapping within the intersections of their interfacial regions required setting weak interactions between chromosomes, we have chosen to set weak fHC-EC attraction to give more freedom for fHC droplets to move through CTs which favor fHC clustering at the interior of the nucleus. Analysis of simulations shows the emergence of different nuclear morphologies governed by the degree of demixing of chromatin types and connectivity of heterochromatin droplets (Fig. 3AB).

**FIG. 3.**
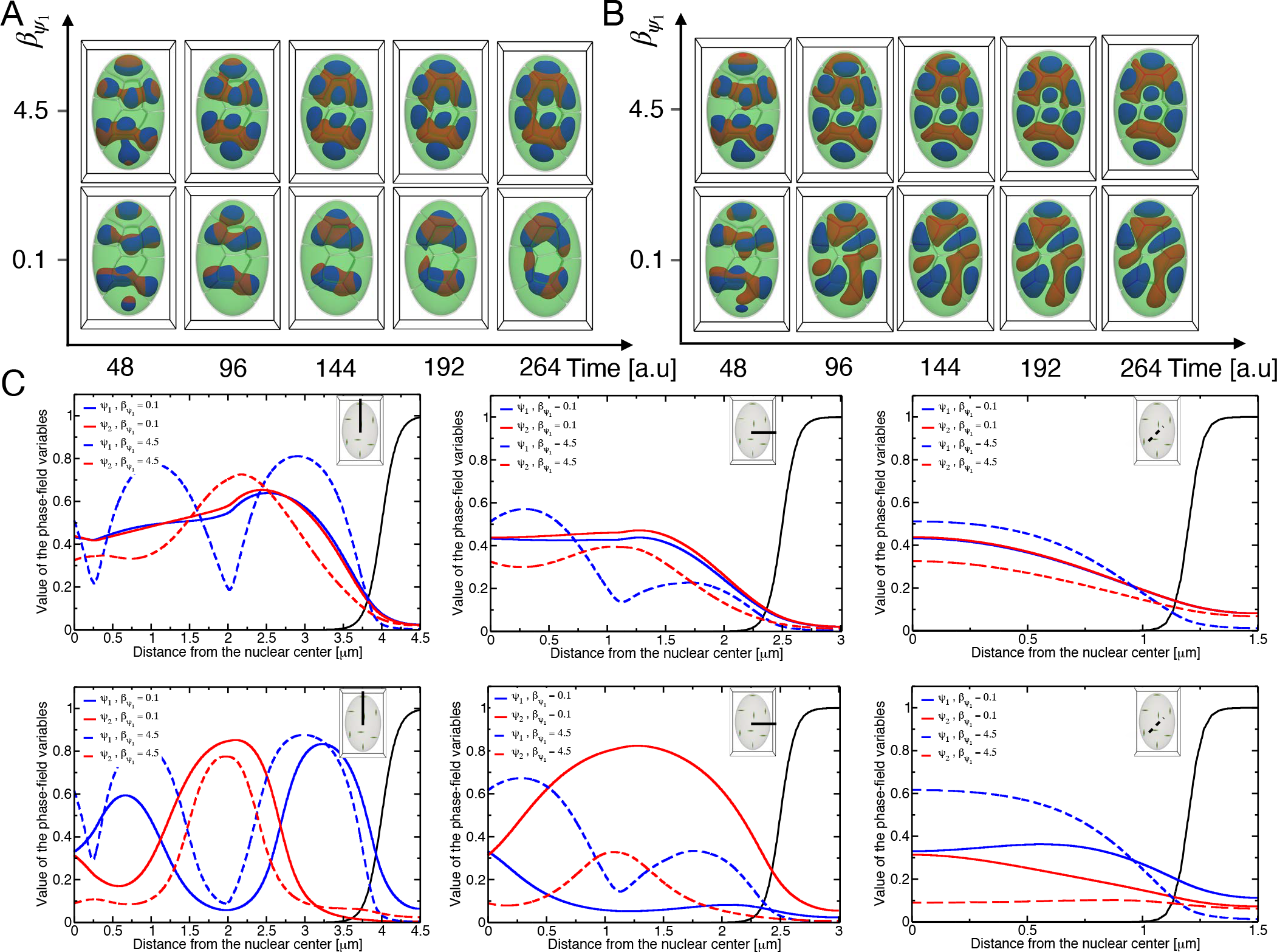
3D chromatin architectures of nucleus resolved with three interacting chromatin types EC (green), fHC (red) and cHC (blue) (A) Nuclear architectures resulting from simulations with constant strong, cohesive interactions between cHC-fHC while varying interaction strength between EC-(cHC, fHC). (B) Nuclear architectures result from simulations with weak cohesive interactions between cHC-fHC while varying interaction strength between EC-(cHC, fHC). (C) Radial density profiles of cHC and fHC, along the major, minor, and *z*-axis. The top panel corresponds to strong, cohesive interactions between cHC-fHC while varying interaction strength between EC-(cHC, fHC). The bottom panel corresponds to weak cohesive interactions between cHC-fHC while varying interaction strength between EC-(cHC, fHC)

Phase-separated HC droplets formation of varying demixed heterochromatin states depending on the strength of interactions between HC types: increasing interaction strength 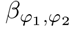 leads to demixing heterochromatin states and forming fully separated HC droplets. On the other hand, lowering the strength of interactions between HC and EC chromatin types within CTs leads to stronger cohesion of HC droplets from neighboring chromosomes that merge into large HC droplets. Simulations show that interactions between HC and EC types of chromatin within individual CT in the lower range values of 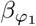and 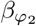are required to establish a physical connection between similar-type of HC droplets, while the distance between the different types of HC droplets increases with the strength of interactions between HC types. As shown, both types of heterochromatin droplets located within CTs display fast motion toward the chromosomal boundaries and clustering in larger HC compartments in the interior of the nucleus for the case 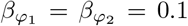. In this case, we noticed a mixing of chromatin types close to the center of the nucleus driven by a strong adhesive interaction between the two HC types. For the case where cHC-EC attraction is stronger than fHC-EC (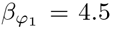 and 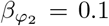), we observed a slow motion of cHC-droplets that restricted their localization at the center of chromosomes, while fHC-droplets moving fast toward the center of the nucleus and clustering in larger compartments. In the remaining cases, we observed demixed chromatin states where the two types of clustered HC are partially or fully disconnected. We shall also notice that the number of fHC clusters within the nucleus is higher than the cHC clusters.

To quantify the effect of multi-component interactions on the spatial organization of chromatin types in the nucleus, we computed the local density profiles of heterochromatin types along the nucleus’s major, minor, and z-axis (Fig. 3 C). The density profiles of both heterochromatin types are similar in the case of strong cHC-fHC attraction compared to EC-(cHC-fHC) interactions, which indicates a mixing state of heterochromatin types residing within the nuclear interior. When the cHC-EC attraction is stronger than fHC-EC, the profiles display demixing states of chromatin of which the formed HC-droplets are close to each other with the separating distance between them depending on the strength of cHC-fHC interactions. The density profiles show that, unlike fHC-droplets, the cHC-droplets reside closer to the nuclear center.

Interestingly, the distance between HC droplets is only slightly influenced by the strength of cHC-EC interactions. Results show that lowering the intensity of cHC-fHC attraction leads to maintaining separated HC droplets. As can be seen, the merged fHC-droplets are partially in contact with the centered cHC-droplets within CTs for strong cHC-EC attraction, while they are fully separated for weak cHC-EC attraction.

To further quantify larger scale spatial arrangement of chromatin compartments, we follow the approach used in [40] based on evaluating spatial descriptors to compute the distance function between similar-type of heterochromatin, volumes, and shapes of cHC and fHC-compartments through sphericity values. We calculated the cumulative distribution function of the distance between the centroid of similar-type HC-droplet positions. We also evaluated the distribution of HC-droplets’ volumes and associated sphericity.

The cumulative distribution functions (Fig. 4) show longer distances between cHC-droplets for simulations generated with the strong attraction between chromatin types cHC-fHC, and EC-cHC. The case with stronger cHC-fHC attraction than EC-cHC shows increased distances between cHC-droplets. One can notice that the distance distribution between fHC-droplets is independent of the strength of cHC-EC interactions for the simulated nucleus generated with strong cHC-fHC attraction. We find the inverse tendency for the distance distribution between HC-droplets in the case with weak cHC-fHC attraction. In this case, we observed more considerable distances between fHC-droplets for strong EC-cHC attraction.

**FIG. 4.**
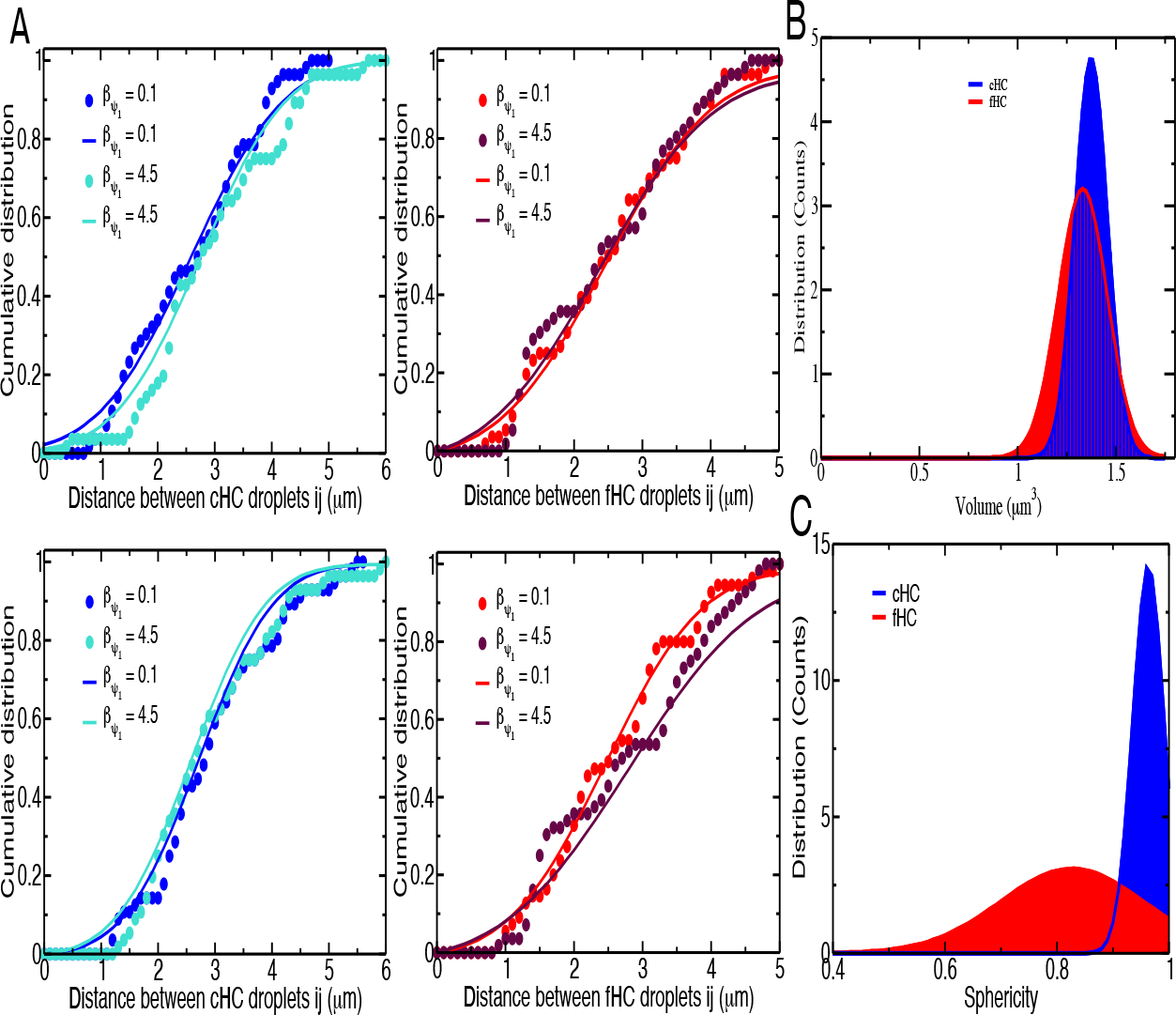
(A) Cumulative distribution of distances between the centroid of *i* −*th* fHC droplets (*i* −*th* cHC droplets) from the *i*-chromosome and the centroids of all remaining *j* −*th* fHC droplets (*j* −*th* cHC droplets) from all *j*-chromosomes (*j* ≠ *i*). The distribution function of the volume of heterochromatin compartments. (C) The distribution function of the sphericity of heterochromatin compartments.

The distribution of volumes of HC-droplets are centered at 1.35*μ m*^3^ and 1.45*μ m*^3^ for fHC and cHC droplets, respectively, and it can be seen that the standard deviation for volume distribution of fHC-droplets is more significant than cHC-droplets (Fig. 4 B). We also noticed the same behavior for the distribution of HC-droplets’ sphericity (Fig. 4C). The distribution of cHC-droplets’ sphericity is centered near the maximal value of 1, which reveals that cHC-droplets adopt a spherical shape. In contrast, the distribution of cHC-droplets sphericity is centered at 0.8, with a significant deviation value compared to the fHC sphericity distribution.

## IV. CONCLUSIONS

The spatial organization of the genome within the eukaryotic nucleus is not arbitrary and plays a pivotal role in gene regulation, replication, and repair. Chromosomes in interphase nuclei are organized into distinct territories (CTs), which are non-randomly positioned with respect to the nuclear periphery and the nucleolus. Studies utilizing fluorescence in situ hybridization (FISH) have delineated these territories, showcasing a radial pattern of genome organization [15, 17]. The radial organization is rationalized by Hi-C analysis using graph theoretical techniques [41, 42] and by polymer modeling of the genome architecture of the nucleus of *Drosophila* at resolution of individual TADs evolving throughout interphase [25]. Eukaryotic nuclei, however, display a dynamic and active organization at all scales, manifesting in the re-organization of epigenetic state and overall architecture throughout interphase, senescence, diseases, and embryonic development [19, 43–46]. Understanding the impact of various equilibrium and non-equilibrium physical forces and biochemical processes on shaping genome organization fuels one of the major unsolved challenges in biophysics known as the four-dimensional nucleosome project [47].

In the present contribution, we developed a multiphase liquid model of the nucleus, which can resolve chromosomal territories, compartments, and nuclear lamina in 4D using a physics-based and data-informed free energy function. Mesoscale Liquid Model of Nuclues (MELON-4D) is a major improvement compared to the 2D mesoscale model we proposed previously [26]. MELON-4D enables rapid hypothesis-driven prototyping of 4D chromatin dynamics, facilitating learning of equilibrium and non-equilibrium driving forces from imaging experiments.

As an application of 4D modeling of eukaryotic nuclei, we explore the interplay of cohesive interactions in chromatin and the adhesive interaction of heterochromatin to the lamina, which influence the dynamics of the radial organization of heterochromatin. We map the phase diagram of nuclear morphologies of *Drosophila*, illuminating the interplay of forces favoring conventional, inverted, and senescent architectures. Interestingly, we have found three distinct morphological phases with different heterochromatin connectivity patterns, which are observed in experiments on different eukaryotic nuclei at different developmental stages [19, 48, 49] suggesting that the same interplay of forces might be underlying these morphologies. In agreement with the previous study of *Drosophila* nucleus [25], we also find that adhesive forces acting between heterochromatin and lamina at the nuclear envelope are the dominant forces that shape chromatin radial organization disruption. Disruption of radial order induced by reduced adhesive interactions leads to the variability of heterochromatin organization.

We have also explored the impact of heterochromatin sub-type interactions on the asymmetry of formed heterochromatin compartments. Namely, we have found that disparity in cohesive interactions between consecutive and facultative heterochromatin could regulate connectivity and asymmetry of heterochromatin compartments without impacting the radial order. Finally, by employing the oblate shape of the nucleus, we quantified asymmetry in radial organization across major and minor axes of the nucleus, highlighting the importance of employing volumetric analysis of nuclear architecture.

We believe that mechanistic modeling of full nuclear dynamics in 4D will experience rapid growth in the future, and understanding the mechanobiology of the eukaryotic nucleus will take center stage in the quest of unraveling the architecture-dynamics-function relation of the genome [50]. We believe MELON-4D model can be a useful framework for physics-based hypothesis testing, which can help rationalize 4D nuclear imaging data.

## Notes

### Competing Interest Statement

The authors have declared no competing interest.

## References

[1] E. de Wit and E. P. Nora, New insights into genome folding by loop extrusion from inducible degron technologies, Nature Reviews Genetics 24, 73 (2023).

[2] M. Conte, E. Irani, A. M. Chiariello, A. Abraham, S. Bianco, A. Esposito, and M. Nicodemi, Loop-extrusion and polymer phase-separation can co-exist at the single-molecule level to shape chromatin folding, Nature Communications 13, 4070 (2022).

[3] L. Mirny and J. Dekker, Mechanisms of chromosome folding and nuclear organization: their interplay and open questions, Cold Spring Harbor Perspectives in Biology 14, a040147 (2022).

[4] S. Brahmachari, V. G. Contessoto, M. Di Pierro, and J. N. Onuchic, Shaping the genome via lengthwise compaction, phase separation, and lamina adhesion, Nucleic Acids Research 50, 4258 (2022).

[5] S. Brahmachari and J. F. Marko, Chromosome disentanglement driven via optimal compaction of loop-extruded brush structures, Proceedings of the National Academy of Sciences 116, 24956 (2019).

[6] E. J. Banigan, W. Tang, A. A. van den Berg, R. R. Stocsits, G. Wutz, H. B. Brandão, G. A. Busslinger, J.-M. Peters, and L. A. Mirny, Transcription shapes 3d chromatin organization by interacting with loop extrusion, Proceedings of the National Academy of Sciences 120, e2210480120 (2023).

[7] G. Spracklin, N. Abdennur, M. Imakaev, N. Chowdhury, S. Pradhan, L. A. Mirny, and J. Dekker, Diverse silent chromatin states modulate genome compartmentalization and loop extrusion barriers, Nature Structural & Molecular Biology 30, 38 (2023).

[8] M. Di Pierro, R. R. Cheng, E. Lieberman Aiden, P. G. Wolynes, and J. N. Onuchic, De novo prediction of human chromosome structures: Epigenetic marking patterns encode genome architecture, Proceedings of the National Academy of Sciences 114, 12126 (2017).

[9] M. Di Pierro, D. A. Potoyan, P. G. Wolynes, and J. N. Onuchic, Anomalous diffusion, spatial coherence, and viscoelasticity from the energy landscape of human chromosomes, Proceedings of the National Academy of Sciences 115, 7753 (2018).

[10] P. S. Carollo and V. Barra, Chromatin epigenetics and nuclear lamina keep the nucleus in shape: Examples from natural and accelerated aging, Biology of the Cell 115, 2200023 (2023).

[11] A. Mahajan, W. Yan, A. Zidovska, D. Saintillan, and M. J. Shelley, Euchromatin activity enhances segregation and compaction of heterochromatin in the cell nucleus, Physical Review X 12, 041033 (2022).

[12] S. T. Jacob, E. M. Sajdel, and H. N. Munro, Different responses of soluble whole nuclear rna polymerase and soluble nucleolar rna polymerase to divalent cations and to inhibition by α-ananitin, Biochemical and biophysical research communications 38, 765 (1970).

[13] I. Eshghi, A. Zidovska, and A. Y. Grosberg, Symmetry-based classification of forces driving chromatin dynamics, Soft Matter 18, 8134 (2022).

[14] I. S. Tolokh, N. A. Kinney, I. V. Sharakhov, and A. V. Onufriev, Strong interactions between highly dynamic lamina-associated domains and the nuclear envelope stabilize the 3D architecture of drosophila interphase chromatin Epigenetics Chromatin 16, 21 (2023).

[15] B. A. Bouwman, N. Crosetto, and M. Bienko, A GC-centered view of 3D genome organization, Curr. Opin. Genet. Dev. 78, 102020 (2023).

[16] P. Das, T. Shen, and R. P. McCord, Inferring chromosome radial organization from Hi-C data, BMC Bioinformatics 21, 511 (2020).

[17] G. Girelli, J. Custodio, T. Kallas, F. Agostini, E. Wernersson, B. Spanjaard, A. Mota, S. Kolbeinsdottir, E. Gelali, N. Crosetto, and M. Bienko, GPSeq reveals the radial organization of chromatin in the cell nucleus, Nat. Biotechnol. 38, 1184 (2020).

[18] N. Crosetto and M. Bienko, Radial organization in the mammalian nucleus, Front. Genet. 11, 33 (2020).

[19] M. Falk, Y. Feodorova, N. Naumova, M. Imakaev, B. R. Lajoie, H. Leonhardt, B. Joffe, J. Dekker, G. Fudenberg, DSolovei, et al., Heterochromatin drives compartmentalization of inverted and conventional nuclei, Nature 570, 395 (2019).

[20] A. Bellanger, J. Madsen-Østerbye, N. M. Galigniana, and P. Collas, Restructuring of lamina-associated domains in senescence and cancer, Cells 11, 1846 (2022).

[21] I. Y. Quiroga, J. H. Ahn, G. G. Wang, and D. Phanstiel, Oncogenic fusion proteins and their role in three-dimensional chromatin structure, phase separation, and cancer, Current Opinion in Genetics & Development 74, 101901 (2022).

[22] K. Vandereyken, A. Sifrim, B. Thienpont, and T. Voet, Methods and applications for single-cell and spatial multi-omics, Nature Reviews Genetics, 1 (2023).

[23] A. M. Rozario, A. Morey, C. Elliott, B. Russ, D. R. Whelan, S. J. Turner, and T. D. Bell, 3d single molecule super-resolution microscopy of whole nuclear lamina, Frontiers in Chemistry 10 (2022).

[24] J. M. Brown, S. De Ornellas, E. Parisi, L. Schermelleh, and V. J. Buckle, Raser-fish: non-denaturing fluorescence in situ hybridization for preservation of three-dimensional interphase chromatin structure, Nature Protocols 17, 1306 (2022).

[25] I. S. Tolokh, N. A. Kinney, I. V. Sharakhov, and A. V. Onufriev, Strong interactions between highly dynamic lamina-associated domains and the nuclear envelope stabilize the 3d architecture of drosophila interphase chromatin, Epigenetics & Chromatin 16, 21 (2023).

[26] R. Laghmach, M. Di Pierro, and D. A. Potoyan, Mesoscale liquid model of chromatin recapitulates nuclear order of eukaryotes, Biophysical Journal 118, 2130 (2020).

[27] R. Laghmach, M. Di Pierro, and D. A. Potoyan, The interplay of chromatin phase separation and lamina interactions in nuclear organization, Biophysical Journal 120, 5005 (2021).

[28] M. Di Bona, M. A. Mancini, D. Mazza, G. Vicidomini, A. Diaspro, and L. Lanzanò, Measuring mobility in chromatin by Intensity-Sorted FCS, Biophys. J. 116, 987 (2019).

[29] R. Bruinsma, A. Y. Grosberg, Y. Rabin, and A. Zidovska, Chromatin hydrodynamics, Biophys. J. 106, 1871 (2014).

[30] D. Saintillan, M. J. Shelley, and A. Zidovska, Extensile motor activity drives coherent motions in a model of interphase chromatin, Proc. Natl. Acad. Sci. USA 115, 11442 (2018).

[31] D. Zwicker, J. Baumgart, S. Redemann, T. Müller-Reichert, A. A. Hyman, and F. Jülicher, Positioning of particles in active droplets, Phys. Rev. Lett. 121, 158102 (2018).

[32] A. E. Slaughter, J. W. Peterson, D. R. Gaston, C. J. Permann, D. Andrs, and J. M. Miller, Continuous integration for concurrent moose framework and application development on github, Journal of Open Research Software 3, (2015).

[33] D. Schwen, L. Aagesen, J. Peterson, and M. Tonks, Rapid multiphase-field model development using a modular free energy based approach with automatic differentiation in moose/marmot, Computational Materials Science 132, 36 (2017).

[34] J. Vazquez, A. S. Belmont, and J. W. Sedat, Multiple regimes of constrained chromosome motion are regulated in the interphase drosophila nucleus, Current Biology 11, 1227 (2001).

[35] S. V. Ulianov, S. A. Doronin, E. E. Khrameeva, P. I. Kos, A. V. Luzhin, S. S. Starikov, A. A. Galitsyna, V. V. Nenasheva, A. A. Ilyin, I. M. Flyamer, et al., Nuclear lamina integrity is required for proper spatial organization of chromatin in drosophila, Nature communications 10, 1176 (2019).

[36] G. Girelli, J. Custodio, T. Kallas, F. Agostini, E. Wernersson, B. Spanjaard, A. Mota, S. Kolbeinsdottir, E. Gelali, N. Crosetto, et al., Gpseq reveals the radial organization of chromatin in the cell nucleus, Nature biotechnology 38, 1184 (2020).

[37] V. O. Chagin, B. Reinhart, A. Becker, O. Mortusewicz, K. L. Jost, A. Rapp, H. Leonhardt, and M. C. Cardoso, Processive dna synthesis is associated with localized decompaction of constitutive heterochromatin at the sites of dna replication and repair, Nucleus 10, 231 (2019).

[38] S. M. Bondarenko and I. V. Sharakhov, Reorganization of the nuclear architecture in the drosophila melanogaster lamin B mutant lacking the CaaX box Nucleus 11, 283 (2020).

[39] P. P. Shah, K. C. Keough, K. Gjoni, G. T. Santini, R. J. Abdill, N. M. Wickramasinghe, C. E. Dundes, A. Karnay, A. Chen, R. E. A. Salomon, P. J. Walsh, S. C. Nguyen, S. Whalen, E. F. Joyce, K. M. Loh, N. Dubois, K. S. Pollard, and R. Jain, An atlas of lamina-associated chromatin across twelve human cell types reveals an intermediate chromatin subtype Genome Biol. 24, 16 (2023).

[40] A. Javier, S. Kaori, G. Valérie, and A. Philippe, Spatial modeling of biological patterns shows multiscale organization of arabidopsis thaliana heterochromatin, Scientific Reports 11, 323 (2021).

[41] P. Das, T. Shen, and R. P. McCord, Inferring chromosome radial organization from hi-c data, BMC bioinformatics 21, 1 (2020).

[42] Y. Zhang, L. Boninsegna, M. Yang, T. Misteli, F. Alber, and J. Ma, Computational methods for analysing multiscale 3d genome organization, Nature Reviews Genetics, 1 (2023).

[43] T. Nozaki, S. Shinkai, S. Ide, K. Higashi, S. Tamura, M. A. Shimazoe, M. Nakagawa, Y. Suzuki, Y. Okada, M. Sasai, et al., Condensed but liquid-like domain organization of active chromatin regions in living human cells, Science Advances 9, eadf1488 (2023).

[44] H. A. Shaban and S. M. Gasser, Dynamic 3d genome reorganization during senescence: defining cell states through chromatin, Cell Death & Differentiation, 1 (2023).

[45] I. Eshghi, J. A. Eaton, and A. Zidovska, Interphase chromatin undergoes a local sol-gel transition upon cell differentiation, Physical review letters 126, 228101 (2021).

[46] H. A. Shaban, R. Barth, L. Recoules, and K. Bystricky, Hi-d: nanoscale mapping of nuclear dynamics in single living cells, Genome biology 21, 95 (2020).

[47] M. Di Stefano, J. Paulsen, D. Jost, and M. A. Marti-Renom, 4d nucleome modeling, Current opinion in genetics & development 67, 25 (2021).

[48] C. L. Smith, Y. Lan, R. Jain, J. A. Epstein, and A. Poleshko, Global chromatin relabeling accompanies spatial inversion of chromatin in rod photoreceptors, Science advances 7, eabj3035 (2021).

[49] E. Miron, R. Oldenkamp, J. M. Brown, D. M. Pinto, C. S. Xu, A. R. Faria, H. A. Shaban, J. D. Rhodes, C. Innocent, S. De Ornellas, et al., Chromatin arranges in chains of mesoscale domains with nanoscale functional topography independent of cohesin, Science advances 6, eaba8811 (2020).

[50] R. Laghmach, M. Di Pierro, and D. Potoyan, A liquid state perspective on dynamics of chromatin compartments, Frontiers in Molecular Biosciences 8, (2022).

